# Effects of Topical Antimicrobial Formulations on *Pseudomonas aeruginosa* Biofilm in an *In vivo* Porcine Burn Wound Model

**DOI:** 10.1101/737338

**Authors:** S.C. Davis, M. Solis, J. Gil, J. Valdes, A. Higa, S. A. Metzger, L. Kalan

## Abstract

Silver has been incorporated into a variety of wound dressings and topical agents to prevent and combat wound infections. *Pseudomonas aeruginosa* is a common cause of burn wound infections and well-known biofilm producer. The objective of this study was to evaluate the effects of a panel of wound dressings containing different silver formulations on *P. aeruginosa* biofilms using an *in vivo* porcine burn wound model. Second-degree burns were created on the skin of specific pathogen-free pigs (n = 3) and inoculated with 2.14 × 10^5^ cfu *P. aeruginosa* per wound. Biofilms were allowed to develop for 24 h, and then each wound was treated with one of 6 treatments: silver oxynitrate dressing (OXY), silver oxynitrate powder (POWD), nanocrystalline silver dressing (NANO), silver chloride dressing (AGCL), silver sulfadiazine (SSD), or a negative control polyurethane film with no silver-based formulation (NEG). Wounds were cultured at D3 post-infection (n = 3 per pig per treatment) and at D6 post-infection (n = 3 per pig per treatment) for quantification of bacteria. On D6, biopsies (n = 3 per treatment) were taken from POWD, SSD, and NEG wounds and wound healing progress was evaluated histologically. At the time of treatment initiation, 24 h post-infection, 8.71 log cfu *P. aeruginosa* were present in burn wounds. On D3 and D6, all treatments significantly reduced bacterial counts in wounds as compared to NEG, but POWD caused an approximately 7-log reduction in bacterial counts on both days and was the only treatment to reduce the bacterial counts to below the threshold for detecting bacteria. The OXY, NANO, and SSD treatments had similar reductions in bacterial recovery on D3 and D6 of approximately 2.5-4 log. The histological healing metrics of reepithelialization percentage, epithelial thickness, white cell infiltration, angiogenesis, and granulation tissue formation were similar among wounds from POWD, SSD, and NEG groups at 6 days post-infection. Silver oxynitrate powder reduced *P. aeruginosa* growth in burn wounds more effectively than other silver-based dressings but did not impact wound healing.

## Introduction

Burn wounds feature a compromised skin barrier that can no longer prevent bacteria from reaching deeper tissues. Burns are especially predisposed to infection due the presence of devitalized tissue and hospital stays that can last for months (Black and Costerton, 2001; Brusselaers et al., 2010; Issler-Fisher et al., 2016). *Pseudomonas aeruginosa* is one species of bacteria commonly isolated from infected burn wounds and is also present in intensive care burn units which often have higher drug resistance rates than *P. aeruginosa* isolated from other wards, even within the same hospital (Yali et al., 2014; Issler-Fisher et al., 2016). *P. aeruginosa* has been reported to be a leading cause of sepsis and death in burn victims (Davis et al., 2008; Brusselaers et al., 2010; Issler-Fisher et al., 2016). Infections with *P. aeruginosa* are of serious concern because this bacterium rapidly produces biofilms, or communities of bacterial cells firmly adhered to a surface and surrounded by an extracellular matrix (Donlan, 2002). Biofilms are difficult to disrupt within wound tissue, in part because biofilms confer increased tolerance to antimicrobial therapy (Black and Costerton, 2001; Davis et al., 2008). Moreover, biofilms can form very quickly; one strain of *P. aeruginosa* isolated from a burn wound produced biofilms *in vitro* within 10 hours (Harrison-Balestra et al., 2003). Therefore, the ideal antimicrobial treatments for burn wounds would have the ability disrupt biofilm produced by pathogenic bacteria.

A survey regarding burn dressings and treatments was sent to healthcare workers from around the world and 84% of respondents believed antimicrobial activity is important in a burn dressing (Selig et al, 2012). This antimicrobial activity will ideally not come from systemic antibiotics, as *P. aeruginosa* is commonly resistant to multiple antibiotics (Potron et al., 2015; Chatterjee et al., 2016) and these drugs must be used prudently to reduce the spread of resistance. Silver is an alternative option to traditional antimicrobials and silver sulfadiazine has been a standard topical treatment for *P. aeruginosa* in burn patients for several decades (Fox, 1968; Barillo and Marx, 2014). In some cases, silver sulfadiazine has been shown to impair the wound healing process (Davis et al., 1990; Mertz et al., 1996; Rosen et al., 2015), thus alternate topical antimicrobial options are still needed. Silver-releasing compounds other than SSD have become popular in wound dressings over the past few decades, and antimicrobial efficacy is often dependent on the formulation of silver used and the delivery mechanism (Meyer et al., 1978; Sullivan et al., 2001). Silver can be coated directly on fibers within wound dressings or can be incorporated into creams that are applied onto burn wounds. Ionic silver in the form of Ag^+^ is essential for antimicrobial activity and is the most common bioactive silver currently available in a wide range of dressing formats. More recently, the novel silver formulation silver oxynitate, that releases the higher oxidation state silver ions Ag^2+^ and Ag^3+^, is hypothesized to have greater bactericidal activity than Ag^+^ because of the higher affinity for electrons, subsequently leading to greater microbial cellular damage (Lemire et al., 2015; Kalan et al., 2017).

The objective of this study was to compare the *in vivo* antimicrobial activity of different silver-formulations incorporated into commercial wound dressings against biofilms of *P. aeruginosa* within porcine burn wounds.

## Materials & Methods

### Experimental Animals

The experimental animal protocol used for this study was approved by the University of Miami Institutional Animal Care and Use Committee, and all procedures followed the federal guidelines for the care and use of laboratory animals (U.S. Department of Health and Human Services, U.S. Department of Agriculture). The studies were conducted in compliance with the University of Miami’s Department of Dermatology and Cutaneous Surgery Standard Operating Procedure. Animals were monitored daily for any signs of pain or discomfort. To minimize possible discomfort, animals were given analgesics throughout the study.

Three animals were used for this study. The young female specific pathogen free (Looper Farms, North Carolina) pigs weighing 40-45 kg were kept in-house for at least 5 days prior to initiating the experiment. The animals were fed a basal diet ad libitum and housed individually in American Association for Accreditation of Laboratory Animal accredited animal facilities with a controlled temperature of 19-21°C a photoperiod of 12 h light and 12 h dark.

### Wounding

For the wounding process, animals were anesthetized, and the paravertebral area of experimental animals were shaved with standard animal clippers. The area was prepared for wounding by washing with a non-antibiotic soap (Neutrogena Soap Bar; Johnson & Johnson, Los Angeles, CA) and sterile water. The skin was then blotted dry with sterile gauze.

Fifty-four second degree burn wounds were made on each animal using 5 specially designed cylindrical brass rods weighing 358 g each that were heated in a boiling water bath to 100°C. A brass rod was removed from the water bath, wiped dry to prevent water droplets from creating a steam burn on the skin, and applied to the skin surface. The rod was held vertically on the skin for 6 seconds, with all pressure supplied by gravity, to make a burn wound 8.5 mm across and 0.8 mm deep. Immediately after burning, the roof of the burn blister was removed with a sterile spatula. Wounds were separated from one another by 6-8 cm of unwounded skin. The day of wounding is D0.

### Wound Inoculation

*Pseudomonas aeruginosa* ATCC 27312, which was originally isolated from a human wound, was used in this study. The challenge inoculum suspension was prepared by scraping overnight growth from a culture plate into 4.5 mL normal saline. This resulted in a suspension concentration of approximately 10^10^ cfu/mL. Serial dilutions were made until a final estimated concentration of 10^6^ cfu/mL was achieved. The final suspension was vortexed and 25 μL was inoculated into each burn wound on D0, the same day the burns were created. All wounds were covered with a polyurethane film dressing (Tegaderm, 3M, St. Paul, MN) within 30 min of inoculation and the dressing was kept in place for 24 h to allow for biofilm formation (Harrison-Balestra et al., 2003). Serial dilutions of the inoculum were plated onto culture media to quantify the viable organisms used for the experiment.

### Treatments

After 24 hours (D1), the polyurethane dressings were removed. Three wounds per treatment per animal were tested for baseline bacterial counts as described in the “Recovery Methods” below. The remaining wounds were randomly assigned to one of six treatment groups: Silver Oxynitrate Dressing (KerraContact Ag) dressing containing silver oxynitrate at 0.4 mg/cm^2^ (OXY; Crawford Healthcare, Doylestown, PA), 150 mg pure silver oxynitrate powder (POWD; Exciton Technologies, Edmonton, AB), 350 mg silver sulfadiazine (SSD; positive control, Watson Laboratories), Nanocrystalline silver dressing (ACTICOAT 7) containing nanocrystalline silver at 1.2-1.4 mg/cm^2^ (NANO; Smith & Nephew, Austin, TX), and a dressing containing AgCl, EDTA, and benzalkonium chloride (AQUACEL Ag EXTRA; AGCL; ConvaTec, Greensboro, NC). For a negative control wounds were treated with polyurethane film dressing only (NEG; negative control). Six wounds per animal were treated with OXY, NANO, or AGCL. Nine wounds per animal were treated with POWD, SSD, or NEG to permit histological analysis.

Each wound was dressed with 5 cm × 5 cm piece of dressing that covered the wounded area and surrounding unwounded skin. Dressings were applied according to the manufacturers’ instructions and dressed wounds from all treatment groups were covered with the NEG polyurethane film dressing to prevent cross contamination. OXY was wet with 200 μL sterile water, which was added on top and then absorbed into the dressing with a sterile spatula. Both POWD and SSD were spread evenly onto the wound with sterile spatulas. NANO was wet with 1000 μL sterile water as per the manufacturer’s directions to activate the dressing properties. AGCL was more absorbent than other treatments and was therefore wet with 4000 μL sterile water. The NEG wounds received no treatment other than the polyurethane film. All wounds were treated on D1, 24 hours after *P. aeruginosa* challenge, and treatments were reapplied on D3, 48 hours after initial treatment application, for final bacterial recovery on D6 with the exception of SSD. SSD was applied every 24 hours as per the manufacturer’s instructions.

### Bacterial Recovery

Three wounds prior to treatment were cultured on D1 to establish the baseline bacterial load within wound. Three wounds per animal per treatment group were recovered on D3 and 3 wounds per animal per treatment were recovered on D6. A sterile surgical steel cylinder with a 22-mm inside diameter was placed around the wound area and 1 mL neutralizer solution was pipetted into the cylinder. The site was scrubbed with a sterile spatula for 30 s and the neutralizer suspension was then serially diluted and plated with the Spiral Plater System (Spiral Biotech, Norwood, MA). The suspension was plated on *Pseudomonas* agar base with cetrimide-nalidixic acid supplement to isolate *P. aeruginosa*. Colonies were counted after 24 h aerobic incubation at 37°C.

### Histology

Three biopsies from each pig were taken from treatments OXY, SSD, and NEG on D6 for histological analysis. Incisional biopsies were obtained through the center of the wounds including normal adjacent skin on both sides. Biopsies were fixed in formalin, stained with hematoxylin and eosin, and sectioned. One section per biopsy was evaluated with light microscopy by a researcher who was blinded to the treatments. Sections were examined for the following elements:

1. Wound epithelialization: the percentage of the wound surface that had been covered with new epithelium.
2. Epithelial thickness: the thickness of the new epithelium within the wound.
3. White cell infiltrate: the amount of subepithelial mixed leukocytic infiltrates. Score: 1 = absent, 2 = mild, 3 = moderate, 4 = marked, 5 = exuberant.
4. Angiogenesis: the amount of new blood vessel formation. Score: 1 = absent, 2 = mild, 3 = moderate, 4 = marked, 5 = exuberant.
5. Granulation tissue formation: the amount of new granulation tissue formation was graded as. Score: 0 = 0, 0.5 = 1-10%, 1 = 11-30%, 2 = 31-50%, 3 = 51-70%, 4 = 71-90%, 5= 91-100.

Digital photographs were also taken of representative wounds at each assessment time to document any signs of erythema and/or infection.

### Statistical analysis

For microbiological analysis, the data were log-transformed and the mean and standard deviation of the log_10_ of the colony forming units/mL (log cfu/mL) were analyzed. Because 1 mL liquid was used to disrupt bacteria from the wounds, the log cfu/mL is used as the measurement for the log cfu recovered from each wound. Log-transformed microbiological results were analyzed using a repeated measures ANOVA in Prism 7 (GraphPad, San Diego, CA). The main effects of time and treatment, as well as the interaction of time and treatment, were tested. Histological data were compared using a Student’s t-test for comparison of all groups. Differences were considered significant when *P* < 0.05.

## Results

### Microbiology

All wounds were inoculated with *P. aeruginosa* on D0. Bacterial biofilm was allowed to develop until D1, at which point the baseline infection was enumerated and all treatments were administered. Silver sulfadiazine is commonly used to control burn infection and was included as a positive control. At D1, 8.71 ± 0.49 log cfu/mL were recovered from the baseline wounds (n = 9; Figure 3A). Treatment group was strongly associated with bacterial recovery at both D3 (*P* < 0.0001) and D6 (*P* < 0.0001) (Figure 3A). However, there was a strong interaction of treatment × time (*P* < 0.0001), as different treatment groups resulted in different reductions in bacterial counts at the two timepoints (Figure 3B). The POWD treatment group had the greatest bacterial reduction as compared to NEG on both days, with a reduction in *P. aeruginosa* counts of 7.17 log on D3 (*P* < 0.0001) and 6.85 log on D6 (*P* < 0.0001). The POWD-treated wounds also had significantly reduced bacterial recovery compared to every other silver formulation on both days (*P* < 0.0001). Bacterial recovery from POWD was below the limit of quantification (LOQ) on D3 and D6 (Figure 3A). On D3, the *P. aeruginosa* yields were similar from OXY, positive control SSD, and NANO, with 2.36, 3.03, and 2.62 log reductions in bacterial recovery compared to the NEG, respectively (*P* < 0.0001; Figure 3B). AGCL caused a reduction in bacterial recovery at D3 of approximately 1.01 log compared to the NEG at D3 (*P* < 0.0001; Figure 3B) but did not decrease recovery as compared to the baseline (*P* = 0.91).

**Figure 1.**
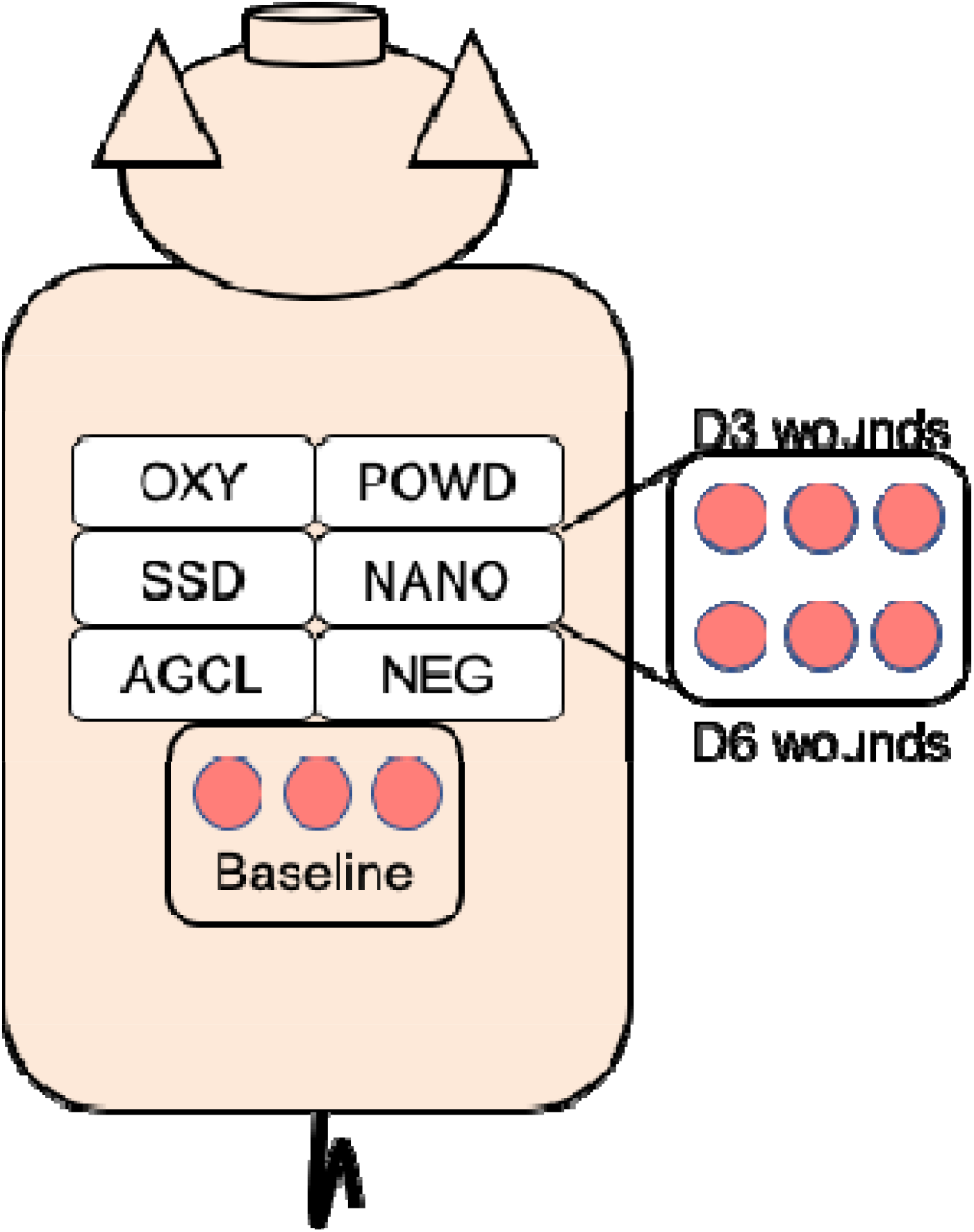
Overview of study design (n = 3 animals). Burns were created on the dorsal side of each animal on D0 and *P. aeruginosa* was administered within wounds at the time of burning. Dressings were applied on D1. Dressings were then removed on D3 for enumeration (n = 3 per treatment per animal) or removed and reapplied for enumeration on D6 (n = 3 per treatment per animal). An additional 3 wounds per animal were created on D0 and biopsied on D6 for the OXY, SSD, and NEG treatments.

**Figure 2.**
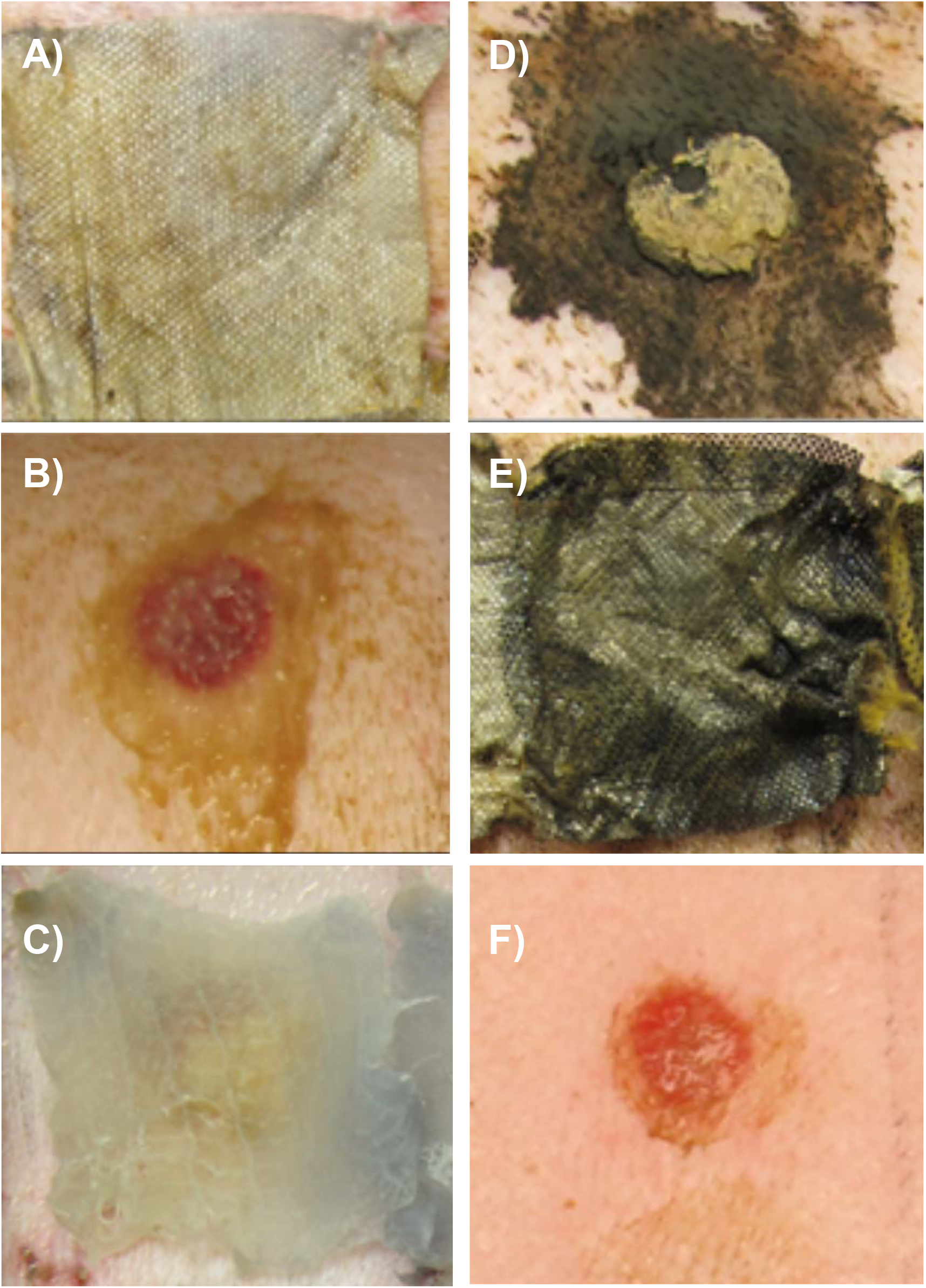
Wound dressings 48 h after application on burns. All dressings were covered with polyurethane film to prevent accidental removal during the experimental period. Additional dressings were as follows: **A)**dressing containing silver oxynitrate at 0.4 mg/cm^2^. **B)**150 mg silver oxynitrate powder. **C**) 1% silver sulfadiazine cream. D) nanocrystalline silver at 1.2-1.4 mg/cm^2^. **E)**dressing containing AgCl, EDTA, and benzalkonium chloride. **F)** no treatment.

**Figure 3.**
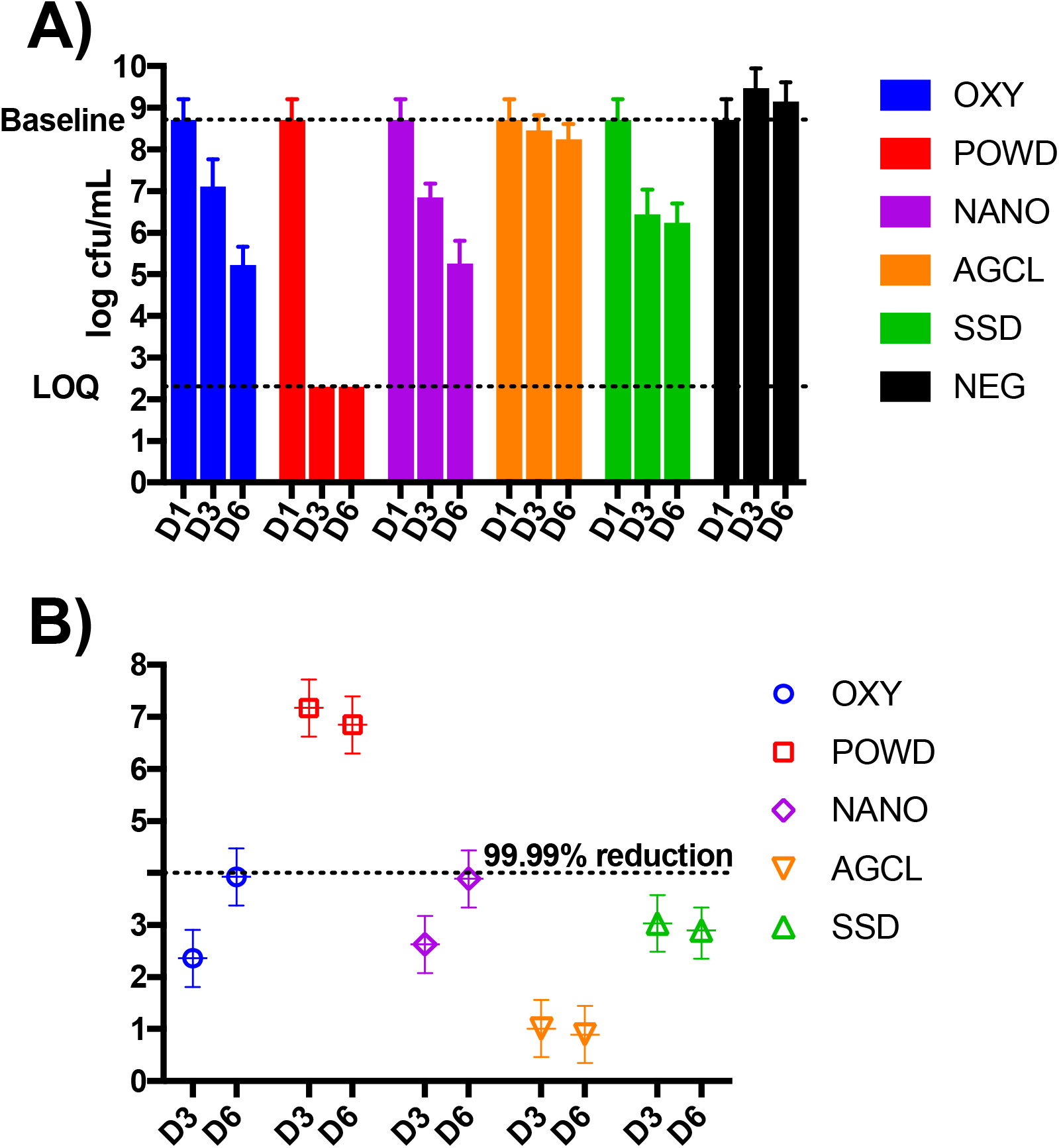
*Pseudomonas aeruginosa* recovery from burn wounds. Burns were created on the backs of pigs (n = 3) and inoculated with 8.59 × 10^6^ cfu *P. aeruginosa* ATCC 27312. Twenty-four h after inoculation, 3 wounds per pig were cultured to establish a baseline bacterial count in the wound. At 24 h post-infection, each treatment (OXY: dressing containing silver oxynitrate at 0.4 mg/cm^2^; POWD: 150 mg silver oxynitrate powder; NANO: dressing nanocrystalline silver at 1.2-mg/cm^2^; AGCL: dressing containing AgCl, EDTA, and benzalkonium chloride; SSD: 350 mg silver sulfadiazine; NEG: polyurethane film with no silver treatment) was applied to 6 burn wounds per pig for microbiological analysis. **A)**Three burns per treatment per pig (9 total burns per treatment) were cultured at D3 post-burn and an additional 3 burn wounds per treatment per pig were cultured at D6 post-burn to measure *P. aeruginosa* recovery. The limit of quantification (LOQ) is the minimum quantifiable bacterial load. **B)**The log reduction in *P. aeruginosa* counts of burn wounds compared to the NEG treatment group.

On D6, the NEG-treated wounds had a log bacterial recovery of 9.15±0.47 cfu/mL. The positive control SSD, OXY, and NANO maintained significant bacterial reductions from D3 compared to the negative control on D6 (*P* < 0.0001). The log reductions compared to the negative control were 2.90, 3.92, and 3.89 for treatments SSD, OXY, and NANO, respectively. The AGCL treatment resulted in a 0.90-log reduction in bacterial recovery relative to the NEG (*P* = 0.001) at D6 (Figure 3B) but were not significantly different from the baseline bacterial recovery (*P* < 0.34).

### Wound appearance

All wounds had slight erythema prior to treatment and this erythema was no longer present by D3. A representative dressed wound from each treatment on D3 after the removal of the polyurethane film is presented in Figure 2A-F. Dressings remained in place for all treatments until being removed by research personnel except for AGCL, which was partially displaced from 3 wounds. The AGCL dressing was slightly adherent to the wound bed during the dressing removal process, one wound from the OXY group had slight adherence to the dressing on D6, and all other treatments were easily removed without adherence to the wound.

### Histology

Three wounds per animal were subject to biopsy and examined histologically at D6 for 3 treatment groups: OXY, SSD, and NEG. The mean epithelialization of the wounds at D6 was 87% (Figure 4A) and mean epithelial thickness was 85 μm (Figure 4B). White cell infiltration was moderate in the 3 treatment groups (Figure 4C) and granulation tissue formation was generally between 11-30% (Figure 4D). Angiogenesis was mild (score 2) for each biopsy, regardless of treatment. None of the 5 histological measures differed among treatment groups.

**Figure 4.**
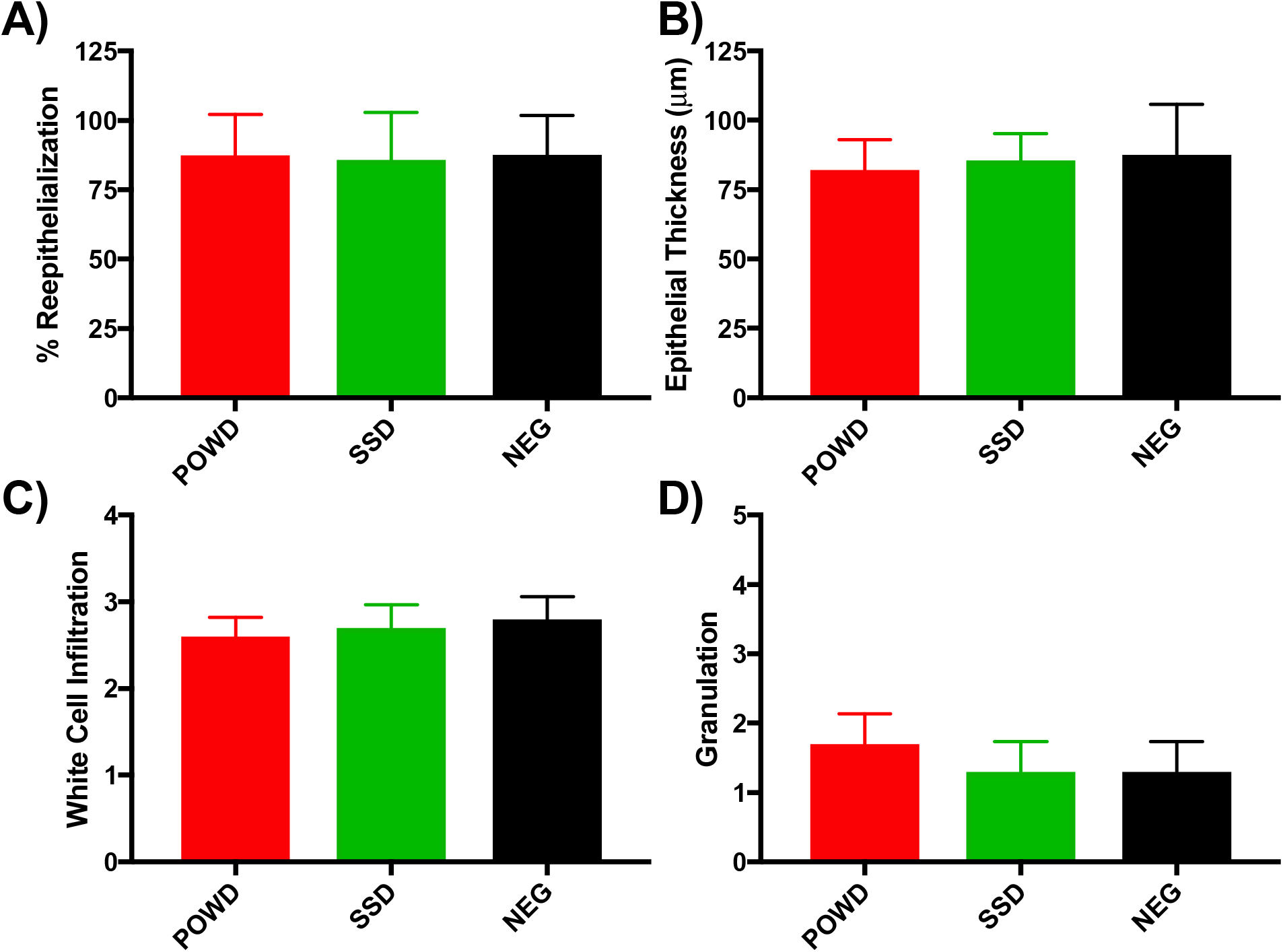
Histological analysis of *Pseudomonas aeruginosa*-infected burn wounds. Burns were created on the backs of pigs (n = 3) and inoculated with 8.59 × 10^6^ cfu *P. aeruginosa* ATCC 27312. At 24 h post-infection, each treatment (POWD: 150 mg silver oxynitrate powder; SSD: 350 mg silver sulfadiazine; NEG: polyurethane film with no silver treatment) was applied to 3 burn wounds per pig for histological analysis. At D6 post-infection, excisional biopsies of burn wounds were collected and one section per biopsy was analyzed histologically by a researcher who was blinded to treatments. **A)**The percent of the burn wound width that was covered with new epithelium. **B)**Thickness of the new epithelial layer. **C)**White cell infiltration within the biopsied wound tissue. Scoring system: 0 = absent, 1 = mild, 2 = moderate, 3 = marked, 4 = exuberant. D) Granulation tissue formation. Scoring system: 0=0, 0.5: 1-10%, 1: 11-30%, 2: 31-50%, 3: 51-70%, 4: 71-90%, 5: >90%

## Discussion

The ideal burn treatment would maintain a moist environment and pH, manage exudate, and prevent infection, all without disturbing the healing process of the burned tissue or causing pain to the patient (Selig et al, 2012). Nearly all respondents to a survey of medical professionals who deal with burn wounds believed that such a dressing does not exist (Selig et al, 2012). Antimicrobial activity in a dressing is critical because infection is a major complication associated with burn wounds (Church et al 2006). Bacteria within burns can form dense biofilms (Kennedy et al 2010) that further impair wound healing. *Pseudomonas aeruginosa*, which can be multidrug-resistant, is a common cause of burn wound infections (Yali et al, 2014; Issler-Fisher et al, 2016; Fournier et al 2016) and well-known biofilm producer. Biofilms produced by *P. aeruginosa* confer antimicrobial resistance in hypoxic regions of the biofilm with reduced bacterial metabolism (Borrellio et al., 2004) or when extracellular DNA chelates cationic peptides (Mulcahy et al, 2008). This biofilm-induced drug resistance, coupled with the high prevalence of antimicrobial drug resistance in *P. aeruginosa* (Potron et al., 2015; Chatterjee et al., 2016), indicates a pressing need to manage burn wound infections without using conventional antimicrobial drugs.

This study aimed to characterize the efficacy of different types of silver formulations commonly used in burn wound management compared to a non-antimicrobial negative control (polyurethane film). Each silver formulation disseminates bactericidal silver ions within the dressing in different ways. Some of the formulations (SSD, NANO, and AGCL) release Ag^+^, while OXY and POWD release silver ions in higher oxidation states (Ag^2+^, Ag^3+^). The higher *in vitro* antimicrobial activity of Ag^2+^ and Ag^3+^ compared to Ag^+^ silver used in antimicrobial dressings is hypothesized to be due to the greater oxidation state of the silver (Lemire et al., 2015), and this higher antimicrobial activity translated into our *in vivo* wounds on porcine skin from the POWD as compared to other treatments. POWD treated wounds resulted in undetectable levels of *P. aeruginosa* from infected burn wounds, translating to greater than a seven-log reduction of viable bacteria.

The United States Food and Drug Administration (FDA) requires a bacterial reduction of at least 4-log (99.99%) by a wound dressing as compared to a negative control treatment for the dressing to be considered antimicrobial (FDA, 2009). Although each silver formulation we tested resulted in a reduced bacterial recovery as compared to the negative control after three and six days of infection, only the silver oxynitrate powder resulted in a log reduction value to fulfill the FDA’s 4-log requirement on both D3 and D6. Even the positive control silver sulfadiazine, which has been in use in burn patients since the 1960s (Barillo and Marx, 2014), only decreased *P. aeruginosa* counts by 3.03 log (99.91%). The AGCL dressing is marketed as an anti-biofilm dressing, however in our study we did not observe significant log reduction of biofilm from burn wounds. While there was a reduction compared to the untreated negative control of ~1 log, we observed no change from the baseline in bacterial counts. This suggests AGCL may suppress further proliferation of *P. aeruginosa* in the tissue, but may only have modest antimicrobial activity against already existing biofilm. Of the traditional contact layer dressings, the OXY and NANO dressings did reached a 4-log level of reduction by D6. The OXY dressing is coated with silver oxynitrate at a concentration of 0.4 mg/cm^2^ resulting in an effective exposure of ~10 mg per dressing in this study. The POWD treatment was administered at a concentration of 150 mg per wound to evaluate potential toxicity at a high dose. Our results suggest that silver oxynitrate is effective at disrupting *P. aeruginosa* biofilms at much lower dose but even at high doses is not toxic.

Burn dressings should have antimicrobial activity without impairing the wound healing process. Silver sulfadiazine’s impairment of wound healing is a well-documented drawback of the treatment (Poon and Burd, 2004; Rosen et al., 2015), and we performed histological analysis for a preliminary comparison of treatments on the wound healing process. Compared to the NEG or SSD, POWD dramatically reduced the number of viable *P. aeruginosa* within the burn wounds without noticeably affecting the speed of the wound healing process. In a recent study, the higher-oxidation state silver released by OXY and POWD resulted in uninfected wounds in mice to close more quickly than wounds that were not treated with silver (Thomason et al, 2018), but the contractile nature of wound closure in mice may translates less directly into human wound closure than the porcine model used in the current study. Detection of differences in wound healing metrics such as epithelialization rate and white cell infiltration would require a greater sample size and longer follow-up time than we were able to utilize. Additionally, the *in vivo* activity of the most effective treatment we used here, the POWD, should be tested in additional burn wound pathogens as silver oxysalts have been demonstrated to have *in vitro* antimicrobial activity against a spectrum of bacteria that includes multi-drug resistant *Acinetobacter baumannii*, *Staphylococcus aureus*, and *Enterococcus faecalis* (Kalan et al., 2017).

## Conclusions

*Pseudomonas aeruginosa* was able to colonize burn wounds on porcine skin and silver-containing treatments reduced bacterial counts in wounds at 2- and 5-days post-infection. Silver oxynitrate powder had the greatest reduction in *P. aeruginosa* biofiom and we were unable to detect changes in wound healing progress compared to the negative control. Silver oxynitrate represents a promising compound with *in vivo* activity for the management of burn wound healing and infection.

## Conflict of Interest Statement

This work was funded by Exciton Technologies. LK was employed by Exciton Technologies at the time the work was completed.

